# Packing of apolar amino acids is not a strong stabilizing force in transmembrane helix dimerization

**DOI:** 10.1101/2025.04.26.649789

**Authors:** Gilbert J. Loiseau, Alessandro Senes

## Abstract

The factors that stabilize the folding and oligomerization of membrane proteins are still not well understood. In particular, it remains unclear how the tight and complementary packing between apolar side chains observed in the core of membrane proteins contributes to their stability. Complementary packing is a necessary feature since packing defects are generally destabilizing for membrane proteins. The question is the extent by which packing of apolar side chains – and the resulting van der Waals interactions – are a sufficient driving force for stabilizing the interaction between transmembrane helices in the absence of hydrogen bonding and polar interactions. We addressed this question with an approach based on high-throughout protein design and the homodimerization of single-pass helices as the model system. We designed hundreds of transmembrane helix dimers mediated by apolar packing in the backbone configurations that are most commonly found in membrane proteins. We assessed the association propensity of the designs in the membrane of *Escherichia coli* and found that they were most often monomeric or, at best, weakly dimeric. Conversely, a set of controls designed in the backbone configuration of the GAS_right_ motif, which is mediated by weak hydrogen bonds, displayed significantly higher dimerization propensity. The data suggest that packing of apolar side chains and van der Waals interactions may be a relatively weak force in driving transmembrane helix dimerization, unless highly optimized. It also confirms that GAS_right_ is a special configuration for achieving stability in membrane proteins.

**Statement of Significance:** The proteins that insert into the cellular membranes provide the cell with essential functionalities. However, how nature designed these proteins to fold into a precise conformation and become active is still poorly understood. In this article we investigate the role of one of the most important factors for protein folding, the van der Waals forces that produce favorable interactions when atoms are in close contact. We know that the formation of extensive van der Waals interactions through contacts are critical for proteins to be stable but we do not know if they are a strong driving force for folding a membrane protein. We addressed this question by designing hundreds of new membrane proteins design based on the optimization of these interatomic contacts and measuring if the resulting proteins come together to form stable pairs. The data suggest that the packing of apolar side chains and the resulting van der Waals interactions are not a strong force for supporting folding in the membrane.

## Introduction

Like all proteins, helical integral membrane proteins need to fold into a specific tridimensional structure to support signaling, transport, catalysis, organization and plasticity and the many other essential functions they perform in the cell membranes. Folding in the membrane is a complex process that requires that proteins find their lowest energy state in an anisotropic environment that spans the hydrophobic core of the membrane, the polar lipid headgroup region and bulk water. Because of this complexity, it has been helpful to subdivide the folding process conceptually into two independent stages, the insertion of the transmembrane helices in the membrane (stage 1) and their association (stage 2) (1, 2). Even though membrane proteins occasionally incorporate within their fold helices that are too short or hydrophilic to be independently stable (3–11), the Two-Stage Model recognizes that most hydrophobic helices of sufficient length to span the membrane are thermodynamically stable in the membrane and can be considered independent folding domains.

The insertion of these helical domains in the membrane (stage 1) is primarily driven by the hydrophobic effect (12–16), i.e., the major driver of the folding of soluble proteins (3, 17, 18). Since this force is spent during insertion and is no longer available for folding, the question becomes what are the factors that promote the association of the helices in the second stage? Polar interactions such as electrostatics, cation→π and, especially, hydrogen bonding can in principle be energetically favorable in the apolar environment of the membrane (19, 20). Indeed, early studies established that individual polar side chains within an otherwise hydrophobic helix are sufficient to drive the self-association of TM helices (21–23). Further experiments suggest that the contributions of individual hydrogen bonds in membrane proteins are relatively modest but not insignificant (between 0.4 and ∼2 kcal/mol) (24–30). Weak hydrogen bonds can also contribute to helix-helix interaction stability, such as those observed in the GAS_right_ interaction motif (31–33), involving activated Cα–H carbon donors (34–37) and backbone carbonyl acceptors on the opposing helix. However, in many cases, the interaction interfaces of TM helical pairs do not display polar interactions. Polar amino acids are relatively rare in TM helices. Furthermore, the most common of them, the mildly polar Ser and Thr, tend to satisfy the hydrogen bonding potential of their hydroxyl group within their local backbone instead of participating in inter-helical hydrogen bonds (38, 39). Therefore, the binding interfaces of helical pairs are often mediated primarily by packing of apolar surfaces and van der Waals (vdW) interactions, raising the question of to what extent these forces can stabilize association.

There is no doubt that packing is a necessary feature of stable membrane proteins. The core of membrane proteins is tightly packed (40, 41) and disruption of packing, either by the creation of voids or the introduction of steric clashes, is generally destabilizing (24, 27, 42–45). Experimental measurements of the cost of creating packing defects within the core of bacteriorhodopsin suggest that the energetic contribution of vdW interactions in membrane proteins (approximately 20 cal mol^-1^ A^2^) is similar to the values obtained for the water-soluble lysozyme (18 cal mol^-1^ A^2^) (46). These studies indicate that packing defects are expensive and that tight packing is necessary for stability but do not demonstrate the extent to which packing is an actual driving force for helix-helix association, beyond being a mere necessity. A library screening analysis showed that TM helices composed of purely hydrophobic amino acids can activate the single-pass PDGFβ tyrosine receptor kinase, suggesting that apolar packing can be sufficient to mediate effective interactions leading to specific biological activity (47). However, the study was based on cellular response and did not confirm if the library elements formed a stable complex with the receptor. To date, the evidence that best supports the case that vdW interactions between apolar side chains can be a significant stabilizing force in membrane protein is a protein design study based on the scaffold of phospholamban, a regulator of the sarcoplasmic reticulum Ca^2+^ pump (48). The pentameric TM complex of phospholamban is mediated by a tightly packed hydrophobic C-terminal region, whereas the N-terminal region is dynamic and contains polar residues. When these polar residues were replaced with apolar side chains, the complex retained its stable pentameric assembly. The authors recognized a steric model based on a repeated LxxIxxx heptad motif (featuring β-branched amino acids at *e* positions and non-branched amino acids at *g* positions) that favored tight packing in the pentameric configuration. These results provide compelling evidence that apolar side-chain packing can be sufficient as a strong driving force for oligomerization in the membrane. However, it was noted that the requirement for complementarity at the interface of this complex was very stringent, to the point that the pentamer can be completely destabilized by minor steric changes. This raises the question of whether the phospholamban model is a highly optimized case and to what extent packing can be a general stabilizing force across a variety of folds.

To answer this question, here we test the propensity of packing to promote association of TM helix dimers. Helical dimers are a simple and tractable system for studying the principles of TM helix-helix association. They are also extremely important biologically. For example, approximately 6% of the human proteome is composed of single-pass receptors, which often activate through ligand-induced stabilization of dimeric conformations (49–51). To understand the role of apolar packing across the variety of dimeric configurations observed in nature, here we use a high-throughput protein design strategy using a large number of starting backbones. We hypothesize is that if steric complementarity and vdW forces are sufficient for driving TM association, it should be possible to use this feature as a principle to design stable dimers.

We compare computationally predicted stability with the experimental dimerization propensity across a series of designed dimers, an approach similar to a previous study in which we explored the energetics of association of the common GAS_right_ dimerization motifs (52). That analysis indicated that vdW interactions and weak Cα–H hydrogen bonds combine to determine the stability of GAS_right_ dimers (52). In the model, the vdW interactions contribute a significant fraction of the predicted binding energetics of these dimers. This suggests that even if the hydrogen bonding component was somehow turned off, the vdW interactions should be sufficient for producing relatively strong association. However, here we found that designs based on packing alone are only weakly dimeric or monomeric. We conclude that, while it may be possible for vdW interactions to produce highly stable structures in some cases, the contributions of packing are either too weak or their requirements are too stringent to be a general strong driving force in the dimerization in TM dimers.

## Results and Discussion

### Design strategy

To assess the general role of packing and vdW interactions in TM helix dimerization, we designed *de novo* a large number of dimers mediated primarily by packing and assessed their propensity to self-associate experimentally. To obtain the most favorable designs, they were based on backbones selected from the most common geometries observed in membrane proteins. The backbone geometries were identified by analyzing all pairs of interacting helices found in membrane proteins of known structure from the Orientations of Proteins in Membranes database, filtered by sequence similarity from the PDB (53, 54). Any two helices in close contact in the structures were considered as an individual helical pair. Since our designs are based on canonical α-helix conformation, unlike previous similar analyses (33, 55), we restricted the pairs to the subset that assumes strictly regular α-helix conformation (excluding all segments that contained kinks, curvature, 3_10_ helix and any other deviations from a standard α-helix) before the conformational parameters of each helical pair were computed (i.e., the crossing angle θ, the inter-helical distance *d*, the axial rotation of each helix ω, and the displacement in along the helical axis or *Z*-shift, Fig. 1a).

**Fig. 1.**
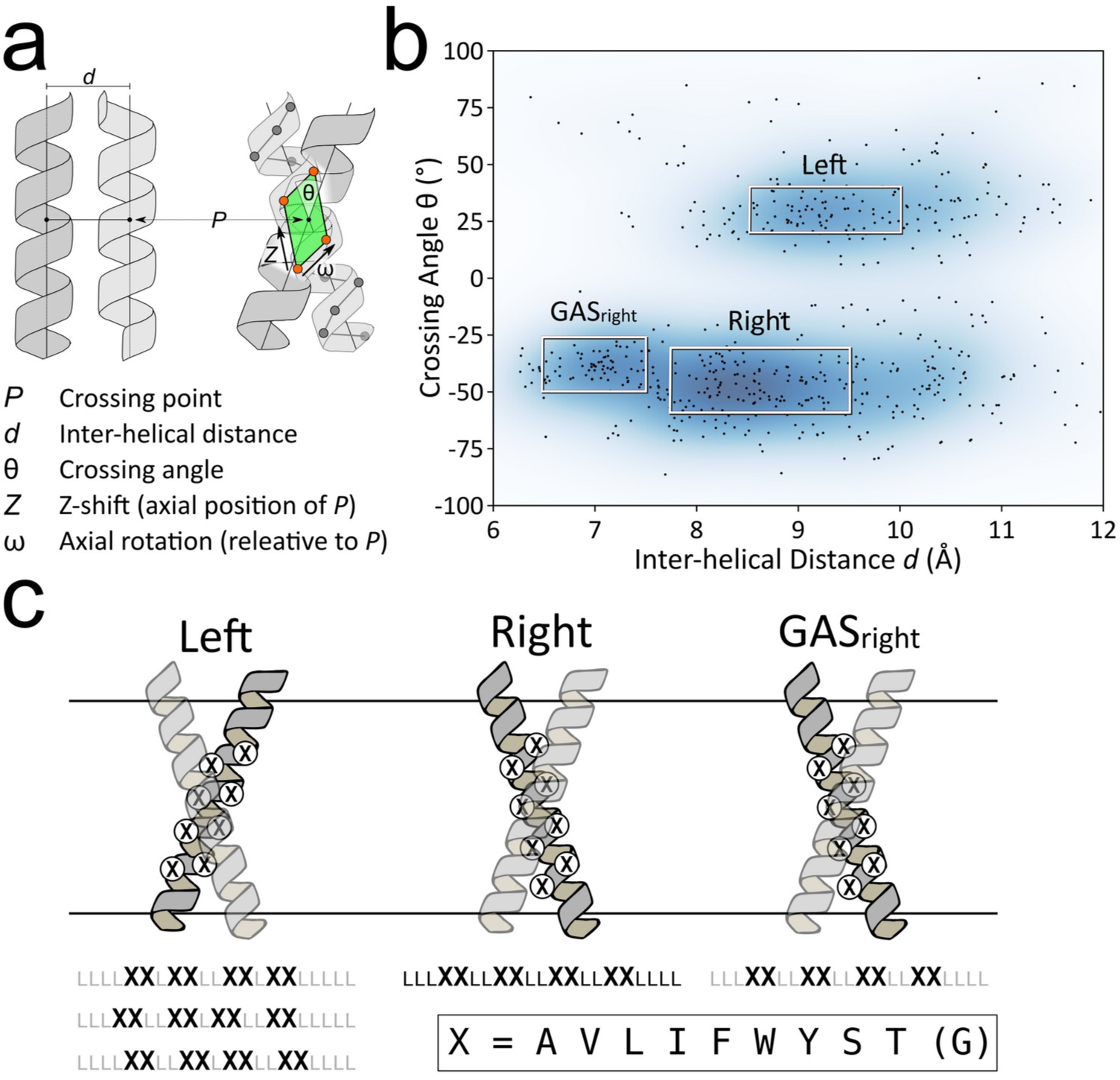
Protein design strategy. a) Inter-helical parameters defining the geometry of a dimer: the inter-helical distance *d*; the crossing angle θ; the rotation of the helix around its axis ω; and the vertical position *Z* of the point of closest approach *P*. The ω and *Z* coordinates are relative to the green parallelogram (the unit cell in the helical lattice) formed by the four Cα of closest approach. b) Distribution of inter-helical distance *d* and crossing angle θ found in pairs of canonical α-helices in membrane proteins of known structure extracted from the Orientations of Proteins in membranes (OPM). The coordinates are overlaid over their kernel density estimation. c) All designs were based on a 21 amino acid backbone, with the crossing point placed in the middle of the membrane, 8 variable interfacial positions (“X”) and all non-interfacial positions set to leucine (“L”). The placement of the interfacial positions was dependent on the ω and *Z* coordinates for designs in the Left region (see supplementary Fig. S1).

Fig. 1b illustrates the resulting helix-helix interaction landscape for parallel helical pairs plotted as a function of inter-helical distance *d* and the crossing angle θ. Two major high-density regions are present. In the left- handed region (positive crossing angle θ), helical pairs are found to interact most frequently with an inter-helical distance between 8.5 to 10 Å of and a crossing angle of 20 to 40°. We call this region the “Left” region (highlighted as a box in Fig. 1b). The right-handed region presents a broader range of populated inter-helical distances, from 6.5 to over 10 Å and crossing angles between -30 and -60°. We selected the maximum density region with inter-helical distance between 7.75 and 9.5 Å and a -30 to -60° crossing angle (named “Right”) as the second source of helical pair backbones.

The section of the right-handed region with the closest inter-helical distance (*d* = 6.5 to 7.5 Å, θ = -25 to -55°) corresponds to the GAS_right_ motif, the well-known helix-helix interaction motif (32, 56). GAS_right_ has a sequence signature of small amino acids (Gly, Ala, Ser) at the dimer interface, forming its characteristic GxxxG and similar sequence motifs (GxxxA, SxxxG, etc.). These small amino acids allow for the short inter-helical distance that brings the helical backbones in contact, which results in the formation of characteristic networks of inter-helical weak hydrogen bonds between activated Cα–H carbon donors and backbone carbonyl acceptor groups on the opposing helix (Cα–H···O=C hydrogen bonds) (33). Since these weak hydrogen bonds contribute to stability in GAS_right_ (52) this region is not considered in our analysis of packing as a driving force. Instead, these designs were used as a positive control to validate the methodology applied to designs in the “Left” and “Right” regions.

Similarly to our previous GAS_right_ study (52), we based our designs on a 21-amino acid poly-Leu backbone with the crossing point between the two monomers placed near the center of the membrane, as illustrated in Fig. 1c. For each dimer, 8 positions at the helix-helix interface were set as variable during the design, whereas all non-interfacial positions were standardized to Leucine, a practice that reduces variability in overall hydrophobicity and helps to obtain relatively consistent expression levels and insertion into the membrane during the experimental phase (supplementary Fig. S2). The right-handed designs (“Right” and “GAS_right_”) follow a typical pattern alternating two variable interfacial positions (**x**) with two fixed positions (L), resulting in a LLL**xx**LL**xx**LL**xx**LL**xx**LLLL sequence (32, 52, 56). In the Left region, the interfacial positions follow the LxxLLxL heptad repeat typical of knobs-into-holes patterns found in leucine zippers (57–59). However, the specific registry of the interface positions within the 21-amino acid helix is dependent on the helical pair coordinates ( *Z*- shift and axial rotation, supplemental Fig. S1). For this reason, three different interfacial patterns, all consisting of 8 variable positions, were adopted for designs in this region, as illustrated in Fig. 1c.

The selection of amino acids used at the variable positions at the interaction interfaces was limited to the most commonly found amino acids in membrane proteins. These included the aliphatic Ala, Val, Leu and Ile, which together represent 43% of the composition of TM helices (60). Also included were the aromatic residues Phe, Trp and Tyr (14% of the composition, in total). Trp and Tyr have a tendency to be enriched in the membrane head-group region and Tyr has a tendency to “snorkel”, and thus expose its η-hydroxyl group to water in the head-group region (61). Finally, we included Ser and Thr, since they are also common in membrane proteins (7% each). These two amino acids contain a hydroxyl group and could potentially form inter-helical hydrogen bonds. However, their side chains tend to satisfy the potential by forming intra-helical hydrogen bonds with carbonyl groups at *i*-3 and *i*-4 (38, 39) and are thus compatible with a helix-helix interaction interface mediated primarily by non-polar interactions. Gly is common in membrane proteins (8%) but it was excluded from the Left and Right designs because it plays a critical role in the interface of the GAS_right_ dimerization motif, to exclude the accidental formation of GAS_right_ structures (Fig. 1b).

The design algorithm started with selecting an initial backbone with crossing angle and inter-helical distance values randomly assigned within the boundaries of each region, as well as random *Z*-shift and axial rotation coordinates. The side chains were selected and optimized with a Monte Carlo procedure, producing the final protein sequence. The structure was further refined by iterative cycles of consisting local backbone moves followed by side-chain optimization (19, 62–65). As in our previous GAS_right_ prediction studies (32, 52), each design was evaluated using a computational energy score composed of a vdW function (66), hydrogen bonding (67), and the IMM1 implicit membrane solvation (68). The procedure resulted in 1045 designed sequences divided across the “Left”, “Right” and “GAS_right_” regions, covering a range of both structure and stability.

To control for the possibility that constructs may associate through different modes than their designed structures, we designed point mutations expected to disrupt association (Fig. 2a). Small interfacial amino acids were changed to the larger Isoleucine residues to create significant clashes within the designed structure (“Clash mutants”). In addition, large amino acids were changed to Ala (”Large→Ala”) to reduce packing at the helix-helix interface. Both Clash and Large→Ala mutants were expected to potentially decrease association, with Clash mutants expected to result in drastic decreases as observed in previous studies (52, 69–73).

**Fig. 2.**
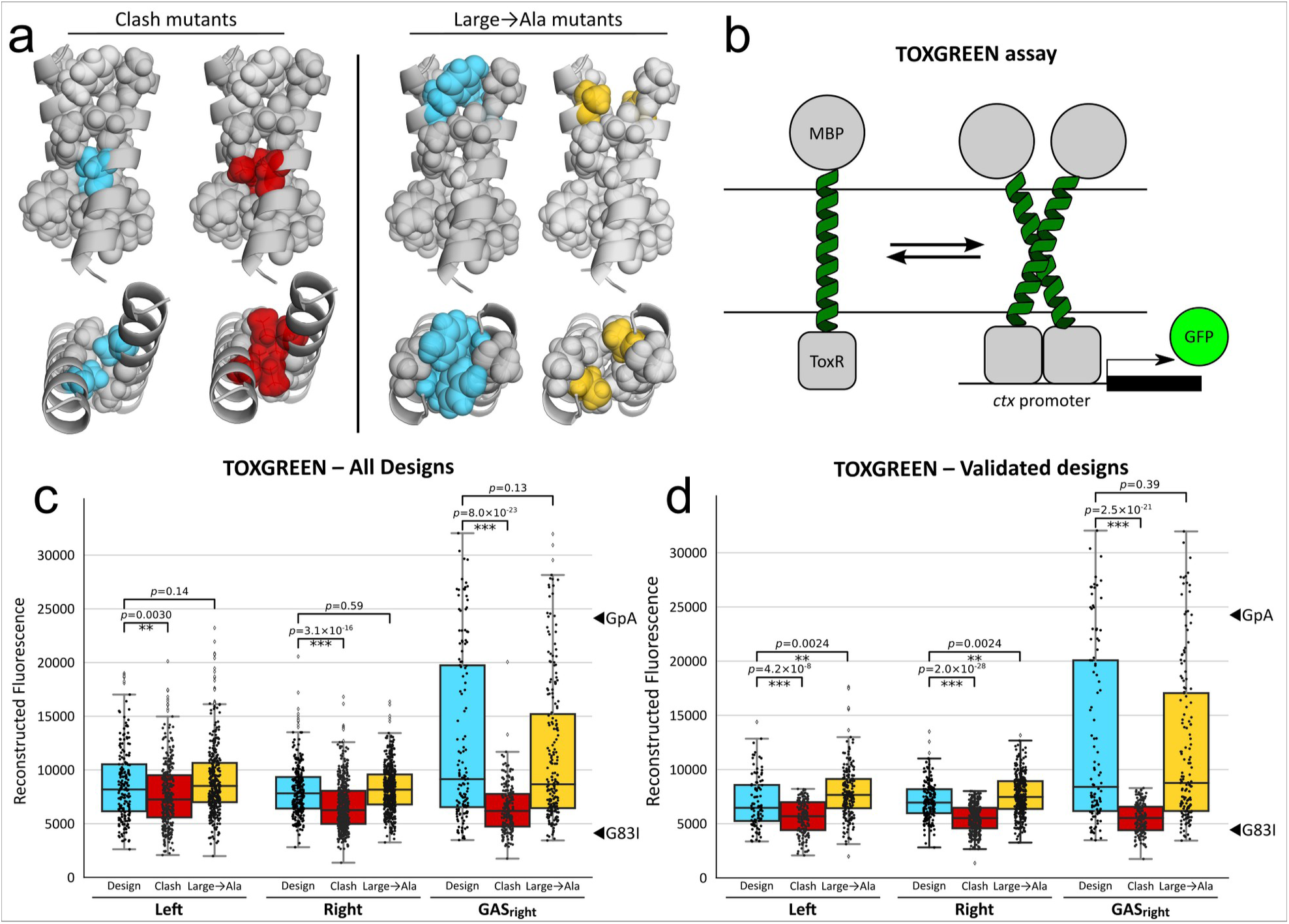
Experimental validation of designed dimers. a) A set of validation mutations was designed for each design. “Clash mutants” are changes to interfacial small amino acids to isoleucine created to introduce significant steric clashes within the designed structure. ”Large→Ala” mutants were created to reduce packing at the helix-helix interface by changing large amino acid to Ala. b) TOXGREEN is an *in vivo* genetic reporter assay that measures oligomerization propensities of TM helices relative to known standards. In the assay, the TM domain under investigation is fused to the ToxR transcriptional activator. TM helix-helix association results in ToxR’s binding to the *ctx* promoter and the consequent in the expression of GFP. In this work, the TOXGREEN assay was used in high-throughput using its sort-seq variant. c) Measurement of dimerization of the designs in the membrane of *E. coli*. The figure displays the reconstructed TOXGREEN sort-seq fluorescence of Left, Right and GAS_right_ designs (blue) and their Clash (red) and Large→Ala mutants. The reconstructed fluorescence of the standards GpA (known strong dimer) and G83I (known monomer) are indicated on the side. The vdW-based designs in the Left and Right regions display a low propensity to dimerize. The GAS_right_ designs include constructs with TOXGREEN signals similar to or higher than the stable GpA standards. The introduction of clash mutations produces a significant reduction of dimerization. d) Same representation as panel c for designs validated by their class mutants (i.e., those designs whose clash mutants are in the monomeric range).

### Experimental strategy

The propensity for dimerization of the designs was experimentally assessed using the high-throughput “TOXGREEN sort-seq” membrane oligomerization assay (74). TOXGREEN (75, 76) is an *in vivo* assay that reports the self-association of TM helices in the biological membrane of *Escherichia coli* (Fig. 2b). The assay is based on a chimeric protein in which the TM domain of interest is fused to the ToxR transcriptional activator. Dimerization of these constructs in the inner membrane induces ToxR binding to a specific promoter, resulting in the expression of GFP. Quantification of fluorescence provides an indication of a TM domain’s propensity for oligomerization. We recently demonstrated that the TOXGREEN signal of a series of GAS_right_ dimers based on a standardized poly-Leu backbone correlates with their thermodynamic stability measured *in vitro* (77). The recently developed high-throughput version of the assay, TOXGREEN *sort-seq,* is suitable for evaluating large libraries of TM helices (74). It is based on Fluorescence-Activated Cell Sorting (FACS), which separates cells in different bins based on GFP expression. The relative frequency of each construct in the various bins is then measured using Next-Generation Sequencing (NGS) and the distribution profile is used to reconstruct the GFP fluorescence of each individual construct.

The direct comparison of the relative dimerization propensity of the various constructs assumes that these are expressed at similar levels in the bacterial membrane. As mentioned in the previous section, the adoption of a standardized poly-Leu backbone substantially reduces expression variability in TOXGREEN (52). The high-throughput workflow of the present analysis does not allow for screening each construct individually and eliminating those that display anomalous expression levels. We verified by Western blot a random subset of 21 constructs, illustrated in supplementary Fig. S2. The majority of the constructs display relatively similar levels of expression. Some variability is expected (observable most notably in constructs R2, R4 and L10, whose bands display reduced density), deriving from inhomogeneity in the rates of protein synthesis, membrane insertion or degradation. While expression variability introduces noise in the apparent dimerization propensities, the large scale of the analysis mitigates this issue, enabling the assessment of statistical trends as long as the majority of the constructs are relatively homogeneous.

### The Left and Right designs are weak in comparison to GAS_right_

The TOXGREEN *sort-seq* analysis recovered 938 of the 1045 (89%) designed sequences. A total of 234 constructs did not grow in maltose minimal media, suggesting that these constructs were inserted poorly in the membrane (76). For 91 designs no validation mutants were found in the reconstruction and were also discarded, resulting in a final set of 613 designs, consisting of 198 Left, 277 Right and 138 GAS_right_ designs (supplemental Fig. S3). The distribution of the computational energy scores of these designs is illustrated in supplemental Fig. S4. The computational energy score of the Right designs was overall less favorable (average -18 kcal/mol) compared to the Left and GAS_right_ (both around -28 kcal/mol), suggesting that the latter backbone configurations are more designable. The structural models of the Right designs also displayed a smaller interface average surface area (median 773 Å), compared to the Left (917 Å) and GAS_right_ designs (858 Å, supplemental Fig. S5a).

Fig. 2c shows the distribution of the reconstructed GFP fluorescence of the designs (blue). The GFP fluorescence values of the standard TOXGREEN controls (the strong dimer glycophorin A and its monomeric G83I variants) are indicated as a reference. We found that the majority of the designs in the Left and Right regions are in a range that indicates that they are monomeric or have a weak propensity to oligomerize. Notably, the GAS _right_ designs span a wider range of fluorescence, including constructs with signals comparable to or higher than the strong GpA dimer. To provide a measure, approximately 40% of the GAS_right_ designs have a reconstructed fluorescence above 60% of GpA (14,500 fluorescent units in the figure), compared to only 8% and 4% of the Left and Right designs. The data confirms the strong propensity of GAS_right_ for self-association and suggests that vdW packing alone is a weak driver for dimerization.

When the “Clash” mutants are considered (red bars in Fig. 2c), a reduction in the distribution of reconstructed fluorescence is observed across all three regions. The reduction is small but statistically significant for the Left and Right designs. A dramatic decrease is observed for the GAS_right_ designs. The data support the hypothesis that a majority of the constructs associate according to their designed interface. The Clash mutants were used to validate a subset of designs that behaved consistently with their predicted interface. This dataset of validated designs included only those “wild type” sequences whose Clash mutants were in the monomeric range using a strict threshold as the criterion (below 35% of the GpA TOXGREEN signal). This selection resulted in the retention of 384 (63%) of the original designs (listed in supplemental Table S1). The distribution of reconstructed TOXGREEN signal of the validated dataset is shown in Fig. 2d. We observed a further reduction of the overall dimerization propensity of the Left and Right designs, indicating that a substantial number of the more stable designs in these regions associated through a spurious interface. The data further confirms a low propensity for the apolar packing designs to self-associate. The distribution of reconstructed fluorescence does not change for the GAS_right_ designs, which retains the entire range of reconstructed fluorescence, including many constructs with fluorescent signals consistent with a strong dimer.

Fig. 2c also shows the reconstructed fluorescence of the sets of Large→Ala mutants (orange) designed to reduce the amount or quality of packing at the interface. Because of the narrow interface of a helix dimer, these mutations do not generally create cavities, which can be energetically expensive (46). Rather, they reduce the interface contact area by approximately 10% on average (supplemental Fig. S5b) and can produce small packing defects. In natural TM dimers, large-to-small mutations can either be well tolerated or reduce dimerization (25, 78–81). For this reason, we hypothesized that we would observe a moderate downward shift in fluorescence distribution. We found that the fluorescence distribution is statistically indistinguishable from WT for the GAS _right_ designs. In the case of the Left and Right designs, we even observed a small fluorescence increase. The data suggest that these mutations designed to reduce packing do not produce an effect measurable with TOXGREEN sort-seq. With regards to the small increase observed in the Left and Right mutants, we speculate that Ala residues may slightly favor non-specific weak association over larger amino acids.

### The energetics of the GAS_right_ designs are consistent with a previous model of association

Comparison of the experimental dimerization propensities with the predicted computational energies can help obtain insights into the energetics of dimerization. Fig. 3a-c show scatter plots of the reconstructed TOXGREEN fluorescence of the designs against the energy scores of their structural models. No notable trend was found for the Left (Fig. 3a) and Right (Fig. 3b) designs. These designs populate only the bottom of the association range of TOXGREEN signal (monomeric or weakly dimeric), therefore it is not possible to assess whether there is any correlation between their association propensity and their computational energetics. Conversely, a statistically significant correlation between association propensity and energy scores is observed for the GAS_right_ designs, a trend that is similar to our previous analysis of association of this motif (52).

**Fig. 3.**
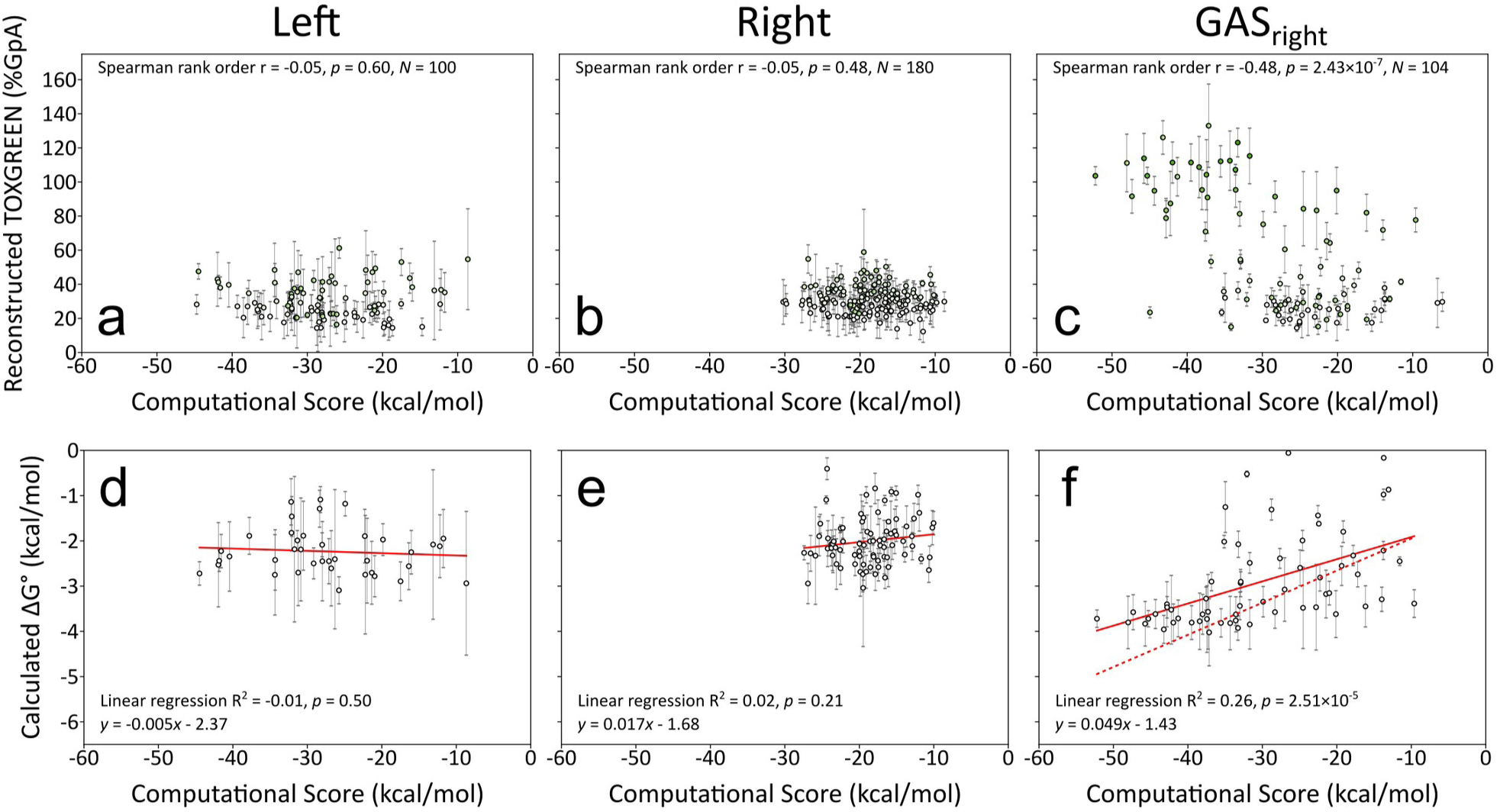
Comparison between computational and extrapolated experimental energetics of the designs. a-c) Comparison between TOXGREEN signal (normalized to the GpA standard) and computational energy score. No trend is observed for the low stability Left and Right designs, whereas a statistically significant relationship is present for the designs with GAS_right_ configuration. d-f) Computational score plotted against an empirical conversion of TOXGREEN signal to ΔG° of association. As in panels a and b, no significant relationship is observed for the Left and Right constructs, which have estimated low dimerization stabilities. A significant linear relationship is observed for the GAS_right_ constructs. The linear relationship (solid line) is relatively similar to the original model of Díaz Vázquez (dashed line).

To further understand the factors that are important for stabilizing the designs, we applied an empirical conversion formula that estimates the free energy of association of constructs from their TOXGREEN signal. This conversion is based on our recent study that found good agreement between the ΔG° of association of a series of poly-Leu GAS_right_ dimers measured *in vitro* in decyl-β-maltopyranoside detergent and their TOXGREEN dimerization propensities (77). Here, we found that there is a statistically significant linear correlation between the computational energy score of the GAS_right_ designs and their calculated Δ*G*° of association (Fig. 3f). The linear regression parameters obtained here (*y* = 0.049*x* – 1.43, solid line, where *y* is the calculated Δ*G*° and *x* is the computational score) are relatively close to the original model of Díaz Vázquez (*y* = 0.076*x* – 1.33, shown in dashed line), indicating that the present set of GAS_right_ designs is consistent with the previous model. As before, no correlation can be found for the weakly associating Left and Right designs (Fig. 3d and e).

## Conclusion

We have presented a large-scale analysis of transmembrane helix association that addressed the question if packing of apolar residues and vdW interactions are an effective driving force for TM helix dimerization. We found that designs based on this feature are monomeric or weakly dimeric in the membrane of *E. coli*. A set of GAS_right_ dimers designed as a control with the same methodology displayed significant propensities to dimerize.

The finding that GAS_right_ dimers are overall more stable than the vdW-based Left and Right designs is not surprising since GAS_right_ and its GxxxG sequence are well-known TM helix interaction motifs that can produce strong dimers. Weak hydrogen bonds contribute significantly to the stability of GAS_right_. However, in our previous energetic model, we estimated that the vdW component represented the largest contributor to association, by approximately a factor of two compared to hydrogen bonding (52). This suggested that even if the inter-helical hydrogen bonding was somehow turned off, vdW packing alone would be sufficient to produce rather stable GAS_right_ dimers. The present data suggest that this is not a general case.

The contributions of vdW interactions in GAS_right_ association may be over-estimated by the model. It is also possible that our high-throughput procedure – which produced thousands of designs across a range of stabilities – may not have had the level of optimization needed for satisfying packing complementarity if these are very stringent. Such stringency was suggested by Mravic and colleagues, who noted that their stable apolar pentameric designs had very strict requirements for complementarity, to the point that the relocation of a single methyl group was sufficient to completely destabilize the oligomer (48). It may be possible that a more intensive computational procedure might identify highly optimized dimers that are stabilized primarily by apolar side-chain packing.

It should also be noted that, even if apolar packing appears much weaker in driving dimerization than interactions involving weak hydrogen bonding, vdW remains a fundamental force in membrane protein folding. First, the association of helical pairs is less costly when they are connected via short loops within the same protein chain; therefore, an interface that produces marginally stable dimers may be sufficiently strong for stabilizing the interaction between two helices in a polytopic membrane protein. Similarly, the association of single-pass membrane proteins often involves extensive interactions between extracellular and intracellular domains, in addition to the TM helices, as, for example, in receptor protein kinases (82); cooperativity with other interacting domains can reduce the energetic requirements for stabilizing TM helix pairs. Additionally, weak interaction can also be important in dynamic biological processes that involve conformational changes, such as the activation and deactivation of signaling molecules (83). Thus, the present study should be viewed as a stringent test for comparing the stability of different interaction motifs and reaffirms the critical role of vdW forces in membrane protein folding.

## Methods

### Structural database analysis of the geometry of ideal helical pairs

A database of non-redundant membrane proteins was obtained from OPM (53) and curated by removing homologous sequences with less than 30% sequence similarity using the program MMseq2 (54). The TM segments were extracted as annotated in OPM and then assessed for their helical nature with a program implemented using the MSL C++ libraries (64). Segments consisting of quadruplets of Cα atoms were assessed with the HELANAL algorithm (84) and were flagged as being helical if their geometry fell within the following parameters: rise per residue: 1.25-1.75Å; rotation per residue: 90-110°; and radius: 2.12-2.42Å. Helical segments composed of at least 13 consecutive helical amino acids were extracted as individual helices. Any two helices with at least 3 Cα carbons within 9 Å of distance from each other were extracted as an individual helical pair. The analysis resulted in a curated database of 835 unique parallel pairs of canonical α-helices extracted from 1541 membrane protein structures. The inter-helical parameters (schematically illustrated in Fig._1a) were then calculated as in Mueller et al. (32), namely the crossing angle θ, the inter-axial helical distance *d*, the axial rotation of each helix ω_1_ and ω_2_, and the position of the crossing point with respect of each helical axis, *Z*_1_ and *Z*_2_. The distribution of the resulting crossing angles and inter-helical distances of the helical pairs are plotted in Fig._1b.

### Computational Dimer Design

Protein design was performed with a program implemented using the MSL C++ libraries on a helical pair consisting of two standard α-helices of 21 amino acids. The backbone coordinates of the pairs were selected by choosing random values for inter-helical distance *d* and a crossing angle θ within the defined regions, as for the following: “Left”: *d* = 8.5 to 10 Å and θ = 20 to 40°; “Right”: *d* = 7.75 to 9.5 Å and θ = -30 to -60°; GAS_right_: *d* = 6.5 to 7.5 Å and θ = -25 to -55°. The axial rotation ω and the *Z* displacement were also assigned randomly in areas of high density. All helical pairs were placed in the middle of a virtual membrane, which was set parallel to the *XY* plane with the *Z* = 0 coordinate corresponding to the center of the bilayer. The *z* coordinate of inter-helical crossing point *P* (Fig. 1a) was placed near this center.

For right-handed designs (GAS_right_ and Right), the variable interfacial positions in the 21-amino acid helix were selected as previously as LLLxxLLxxLLxxLLxxLLLL (52), where x corresponds to a variable position and L corresponds to a non-interfacial position, which was set to Leucine. In order to identify the interfacial position of the left-handed backbones, we calculated the steric requirement as a function of the four geometric parameters *d*, θ, ω and *Z.* As shown in supplemental Fig. S1a, the analysis identified 3 distinct “stripes” of configurations compatible with knobs-into-holes packing. The first stripe ranged approximately from ω=0° and *Z*=1 Å to ω=40° and *Z*=6 Å, the second from ω=40° and *Z*=0 Å to ω=80° and *Z*=6 Å, and the third covered approximately from ω=80° and *Z*=0 Å to ω=100° and *Z*=1 Å. These three regions produced different sets of interfacial residues, corresponding to variable positions patterns LLLxxLLxxLxxLLxxL, LLLLxxLxxLLxxLxxL and LLLLxxLLxxLxxLLxxL, respectively, as illustrated in supplemental Fig. S1b.

Designs were performed using a Monte Carlo sequence optimization algorithm implemented with MSL, starting from 5000 backbones from each of the Left, Right and GAS_right_ regions. The interfacial positions were allowed to assume any of the amino acids most prevalently found in membrane protein sequences, Ala, Val, Leu, Ile, Phe, Tyr, Trp, Ser and Thr. Gly was also allowed for designs based on GAS_right_ geometry. The side chain mobility was modeled using the Energy-Based Conformer Library applied at the 95% level (85). Non-interfacial positions were restricted to Leu and sampled at the 60% level. The variable positions were initially set to a random sequence and then the sequence was optimized with Monte Carlo moves consisting of randomly changing the amino acid at one position, followed by side chain optimization performed with 3 cycles of a greedy algorithm (86). The initial search consisted of 50 cycles from a temperature of 5000 K and cooled to 0.5 K an exponential cooling schedule. Once the end temperature is reached, another search consisting of 100 cycles at a temperature of 3649 K was used to further explore local sequence variation.

Energies were calculated using the CHARMM22 van der Waals function (66), the IMM1 membrane implicit solvation term (68), and the hydrogen bonding function of SCWRL4 (67) as implemented in MSL (64). For Cα–H hydrogen bonds the following parameters were used, as reported previously: *B* = 60.278, *D*_0_ = 2.3 Å, σ*_d_* = 1.202 Å, α_max_ = 74.0°, and β_max_ = 98.0° (32). The dimer energy was calculated as a binding energy by subtracting the energy of the monomer. During the search, the monomeric energy was approximated by a baseline monomer energy calculated from the identity of each amino acid at each position. Additionally, a sequence entropy term was added to penalize low-complexity sequences. This term is sufficiently strong to steer the overall composition of the pool of constructs to approach that of natural membrane proteins, while allowing each individual dimer to adopt the sequence most favored by the interaction energetics.

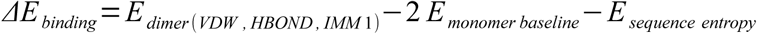

The baseline and sequence entropy terms were not used to compute the final binding energy of a design. Instead, the energy of a side chain-optimized monomer was subtracted from the energy of the dimer.

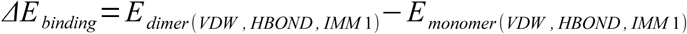

### Monomer energy approximation with baseline energies

A monomer energy approximation was developed by measuring the interactions (van der Waals, hydrogen bonding and solvation energies) for individual amino acids in a large set of monomeric helices of different amino acid compositions. The baseline energy includes a self term (interactions within the single amino acid) and pair term (interaction of the amino acid with other amino acids in the helix), which were computed for 10000 random sequences in a helical configuration and averaged. The pair terms are limited to 10 positions following an amino acid in the helix.

During design, for a given sequence, its monomeric baseline energy *E_baseline_* was calculated as:

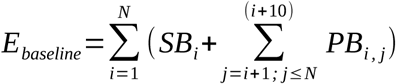

where *i* and *j* are the positions in a helix, *N* is the total number of amino acids in the helix, SB*_i_* is the self baseline energy at position *i*, and *PB_i,j_* is the pair baseline energy between positions *i* and *j*. SB*_i_* is dependent only on the identify of the amino acid at position *i* whereas *PB_i,j_*is dependent on the identities of positions *i* and *j* and their distance in the primary sequence *d*=*j*-*i*.

Supplemental Fig. S6 displays the correlation between computed monomer energy and their baseline monomer energy estimates. The table of self and pair baseline energies is provided as supplemental files Baselines.xlsx.

### Sequence Entropy Term

The sequence entropy term was calculated from the probability of each amino acid in the transmembrane domains extracted from the OPM database.

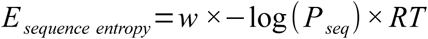

where *w* is a weight factor of 10, *P_seq_* is the probability of a given sequence, R is the gas constant and T is temperature defaulted to 298 K. *P_seq_* was calculated as

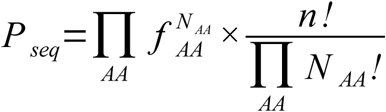

where *N_AA_* is the number of amino acids of type AA found in the sequence and *f_AA_* is the frequency of the amino acid in the OPM database, reported in supplemental Table S2.

### Final structure refinement

After each design was completed, its structure was subjected to a final Monte Carlo procedure to optimize the backbone for the final sequence. The procedure consisted of 100 cycles, with an initial temperature of 1000K cooled to 0 K with an exponential schedule, where the four geometric parameters (*d*, θ, ω and *Z)* were locally varied and the side chains were optimized with 10 cycles of a greedy optimization algorithm (86). This relaxation was repeated 3 times for each protein. The final energy after this backbone relaxation step was subtracted from the energy of a side chain-optimized monomeric helix with the same sequence and used as the design computational energy score.

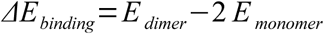

### Library cloning

DNA sequences for the designs were ordered as an oligo pool from Twist Biosciences. Individual segments were amplified using primer pairs specific to each segment by quantitative PCR (Roche KAPA SYBR Fast) based on the protocol outlined in Kosuri et al. (87) followed by traditional PCR reaction using the qPCR reaction as a template. Segments were digested using restriction enzymes NheI-HF and DpnII. The pccGFPKan (TOXGREEN) vector (75) was digested in preparation for ligation using restriction enzymes NheI-HF, BamHI-HF, and CIP. Plasmids were assembled by incubating segment DNA with vector in a 1:10 backbone:insert ratio with ElectroLigase for 2 hours at room temperature, followed by 10 minutes at 65°C to inactivate the enzyme. The inserts were designed such that the last two nucleotides were “TT” before the DpnII cut site “G” resulting in coding an in-frame Leu once ligated into the corresponding pccGFPKan restriction sites resulting in the following general insert sequence RAS**LLLxxLLxxLLxxLLxxLLLL**LILI. 5 μL of the ligation mixture were added to 50 μL of NEB 10-beta *E. coli* cells and cloned through electroporation (BIO-RAD MicroPulser Electroporator), grown for 1 hour in 950 μL NEB Stable Outgrowth Medium media in 37 °C shaker, and diluted 1 mL in 4 mL LB media containing 100 μg/mL ampicillin. 50 μL of a 1:10 and 1:100 dilution were plated on LB plates containing 100 μg/mL ampicillin. Plates were grown for 14-16 hours at 37°C and colony forming units (CFUs) were calculated for each segment (number of Colonies × dilution/number of sequences in the segment). A CFU > 5 was accepted. Overnight cultures corresponding to plates were spun down and the plasmid DNA was extracted (Qiagen). The plasmid library was transformed through electroporation into 50 μL malE deficient *E. coli* strain MM39, grown for 1 hour in 950 μL SOC media in a 37°C shaker, and 100 μL of 1:10 dilution plated on LB plates containing 100 μg/mL ampicillin with the rest being grown overnight in 3 mL LB media containing 100 μg/mL ampicillin. CFUs were again counted and any plates with values > 5 were accepted. Overnight cultures for each segment were stored in 25% glycerol stocks. 50 μL samples and controls from glycerol stocks were grown in 3 mL LB media containing 100 μg/mL ampicillin for 2-4 hours to reach an optical density of approximately 0.1 OD _600_ and the corresponding number of sequences per sample were used to calculate how much to add to a full library glycerol stock. All enzymes acquired from NEB.

### TOXGREEN sort-seq

TOXGREEN sort-seq was performed according to Anderson et al. (74). 50 μL of library glycerol stock were grown in 3 mL LB containing 100 μg/mL ampicillin overnight for 14-16 hours in a 37 °C shaker. FACS was performed using a Sony MA900 cell sorter. The FSC threshold was set at 0.05%. The sensor gains for FL1 (GFP) and backscatter (BSC) were set at 90% and 30%, respectively. The first gate was drawn to select for cells by eliminating cell debris on a side scatter area (SSC-A) vs. forward scatter area (FSC-A) plot. The second gate was drawn to eliminate doublet cells on a forward scatter height (FSC-H) vs. FSC-A plot. 10 μL of sample culture were diluted in 50 mL PBS buffer and 10,000 events were recorded per second in the Fluorescence-Activated Cell Sorter. Control cells containing the GpA, G83I, and No-TM controls were flowed to calibrate the instrument and determine proper gating to remove dead cells. Individual sample libraries were first flowed through the instrument and the fluorescence profile was subdivided into 4 bins. 100,000 events were sorted into bins 1-3, and 50,000 events were sorted into bin 4. Each library sample was sorted in triplicate from biological replicates of overnights into 2 mL LB broth containing 100 μg/mL ampicillin. Sorted populations were grown for 1-5 hours to reach an optical density of approximately 0.3 OD_600_ and plasmid DNA was extracted. Samples were prepared for NGS using PCR: each sorted population was amplified with unique stem primers, allowing us to distinguish between populations for each NGS run. These amplified samples were sent to the DNA Sequencing Center at the University of Wisconsin-Madison Biotech Center for library preparation and next-generation sequencing for 500 reads for each sequence in the library. Library Preparation Services: Index PCR (TruSeq). Sequencing Services: Illumina (NovaSeq) Sequencing [2x150 Shared (10M read increments)].

The GFP fluorescence of each individual construct in the library was reconstructed as in Anderson et al. (74) using a weighted average approach (88). This method normalizes the NGS reads per construct per bin with the fraction of the population found in that bin. To ensure NGS data quality, a cutoff of 1 incorrect base pair per sequence was implemented based on the Phred score (Q) of each base, measuring the probability that the base is incorrect where Q = -10 log_10_P (89, 90). Further, a minimum cutoff of 10 copies of a particular sequence had to be present in the NGS for it to be retained. The normalized fractional contribution *a_ij_*for each construct *i* in each bin *j* was calculated as

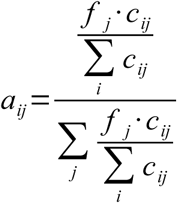

where *c_ij_* is the NGS sequencing count for construct *i* in bin *j*, and *f_j_* is the fraction of the sorted sample population in bin *j*. The reconstructed fluorescence (*RFU_i_*) for each individual protein sequence, expressed as Relative Fluorescence Units was then derived by summing the product of the normalized fractional contribution *a_ij_* and the median fluorescence *m_j_* over each sorted bin *j*.

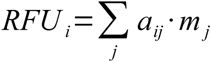

### Normalized TOXGREEN conversion

TOXGREEN fluorescence is typically converted to a relative signal by normalizing its fluorescence to the fluorescence of the strong glycophorin A (GpA) dimer. To reduce the dependency of the normalization from the variability associated with an individual construct, we created a calibration set of approximately 10 constructs to be used for this normalization. Standard TOXGREEN was performed on a set of individual constructs from each design region, including GpA and its monomeric variant G83I, used as the negative control. The fluorescence values, the normalized (% GpA) values and the reconstructed fluorescence values of these constructs are reported in supplemental Table S3. As illustrated in Fig. S7, linear regression was used to determine the relationship between the individual TOXGREEN signal of each construct, expressed as % GpA, and their reconstructed fluorescence. The resulting linear regression equation was then applied to transform the reconstructed fluorescence values to % GpA normalized values.

### Conversion of TOXGREEN signal to an estimated free energy of association

TOXGREEN data was converted to an estimated ΔG of association in detergent using a previously found relationship (77). First, a conversion factor was applied to account for the difference between TOXGREEN and TOXCAT signals.

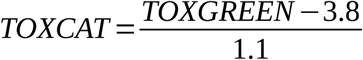

Then a theoretical fraction dimer *FD* was calculated according to the following relationship (77):

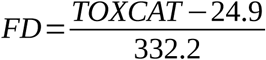

The fraction dimer was converted to a dissociation constant *K_D_*, assuming a concentration of protein χ*_T_* of 1.8×10^-4^ protein molecules per detergent molecule.

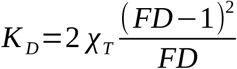

Finally, an estimated free energy of association in detergent Δ*G* was calculated from the dissociation constant, where R is the gas constant and T is 4 °C.

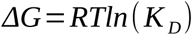

### Immunoblotting

Protein expression was confirmed for an assortment of designs and mutants using immunoblotting (supplemental Fig. S2). Cell lysates were normalized by ThermoScientific Pierce Protein BCA Assay Kit and 10uL loaded onto NuPage 4-12% bis-tris SDS-PAGE gels (ThermoFisher) and then transferred to PVDF membranes (VWR) for 1 hour at 100 millivolts. Blots were blocked using 5% milk (Nestle Instant Nonfat Dry Milk) in TBS-Tween buffer (50 mM Tris, 150 mM NaCl, 0.05% Tween 20) for 2 hours at 4 °C and incubated overnight with peroxidase-conjugated monoclonal anti-maltose binding protein antibodies (Sigma-Aldrich). Blots were developed with Pierce ECL Western Blotting Substrate Kit; 1 mL of ECL solution was added to the blot and incubated for 90 seconds. Chemiluminescence was measured using an ImageQuant LAS 4000 (GE Healthsciences).

### MalE Complementation Assay

To confirm proper membrane insertion and orientation of the TOXGREEN constructs, the library was tested with a MalE Complementation Assay performed in M9 media with maltose as the sole carbon source. The MM39 *E. coli* strain used in the TOXGREEN assay is Maltose Binding Protein (MBP)-deficient. TOXGREEN constructs can support growth in M9 maltose minimal media only if the chimeric protein is expressed and inserted into the inner membrane with the C-terminal MBP moiety correctly oriented to the periplasmic side (76).

The library version of the MalE Complementation Assay in liquid media was performed as described previously (74). Briefly, overnight cultures of each library were used to inoculate 500 mL M9 media containing 0.8% maltose (w/v) with a dilution to a final cell concentration of 0.00125 OD_600_. Samples were taken at time points 30 and 36 hours in triplicate of growth at 37°C. DNA was extracted and submitted for NGS analysis. Samples from the overnight LB cultures used for initial inoculation were also analyzed by NGS for comparison with non-selective conditions. All sequences that had a fractional NGS count change worse than a set of known maltose-negative (non-inserting) controls (constructs “1E01”, “1E11”, “2E11”, “2F12”, “2H11”, “1F02”, “2H07”) (74) were excluded from our analysis.

### Programs and data availability

Programs used to determine the preferred interhelical geometries from the structural database and design the dimers are available at the GitHub repository https://github.com/TmdimerDesign/vdwDesign/, also available at https://doi.org/10.5281/zenodo.16876329. TOXGREEN plasmids for control constructs are available in the AddGene repository (https://www.addgene.org). These include the original TOXGREEN plasmid (pccGFPKAN: AddGene ID #73649), the TOXGREEN positive (GpA: #73651) and negative controls (G83I: #73650) (75) and an extended set of maltose-negative and maltose-positive control constructs (1E01: #239013; 1E11: #239083; 2E11: #239084; 2F12: #239085; 2H11: #239086; 1G01: #239087; 2H01 #239760; 2H07: #239088) (74). The PDB files of the designed dimers are available for download from the Dryad repository at https://doi.org/10.5061/dryad.5dv41nsjg. The repository also include raw sort-seq data used for the calculation of TOXGREEN sort-seq values and for the MalE Complementation Assay. Scripts for the analysis of TOXGREEN sort-seq data are available at https://doi.org/10.5061/dryad.qnk98sfv8 (74).

## Author Contributions

**GJL**: Conception and Design, Data Acquisition, Analysis, and Interpretation, Wrote Manuscript.

**AS**: Conception and Design, Data Analysis, and Interpretation, Wrote Manuscript.

## Acknowledgments

This work was supported by National Institutes of Health grant R35-GM130339 to A.S. and National Science Foundation grant CHE-1710182 to A.S.. G.J.L. acknowledges the support of National Institutes of Health training grant T32GM008505 to the Chemistry-Biology Interface Training Program (CBI).

## SUPPLEMENTARY INFORMATION

**Fig. S1.**
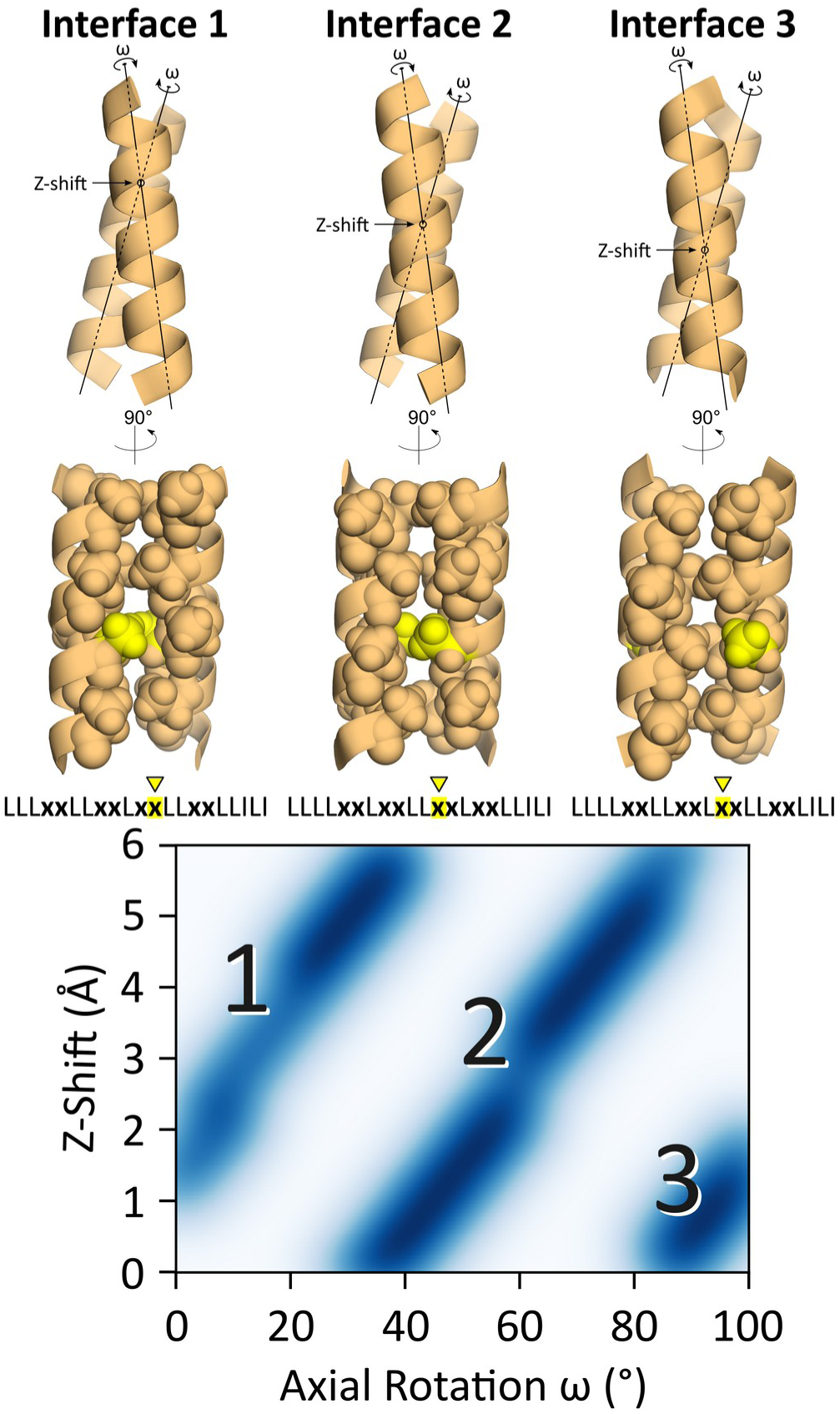
Selection of the interfacial position for the “Left” designs. The interaction interface in the 21-amino acid TOXGREEN construct for the “Left” designs depends on the ω (axial rotation) and *Z*-shift (the position of the crossing point relative to the helical axis. These two coordinates generate three different regions of know-into-hole packing, corresponding to three different arrangements of the variable (**x**) interfacial position during the design process.

**Fig. S2.**
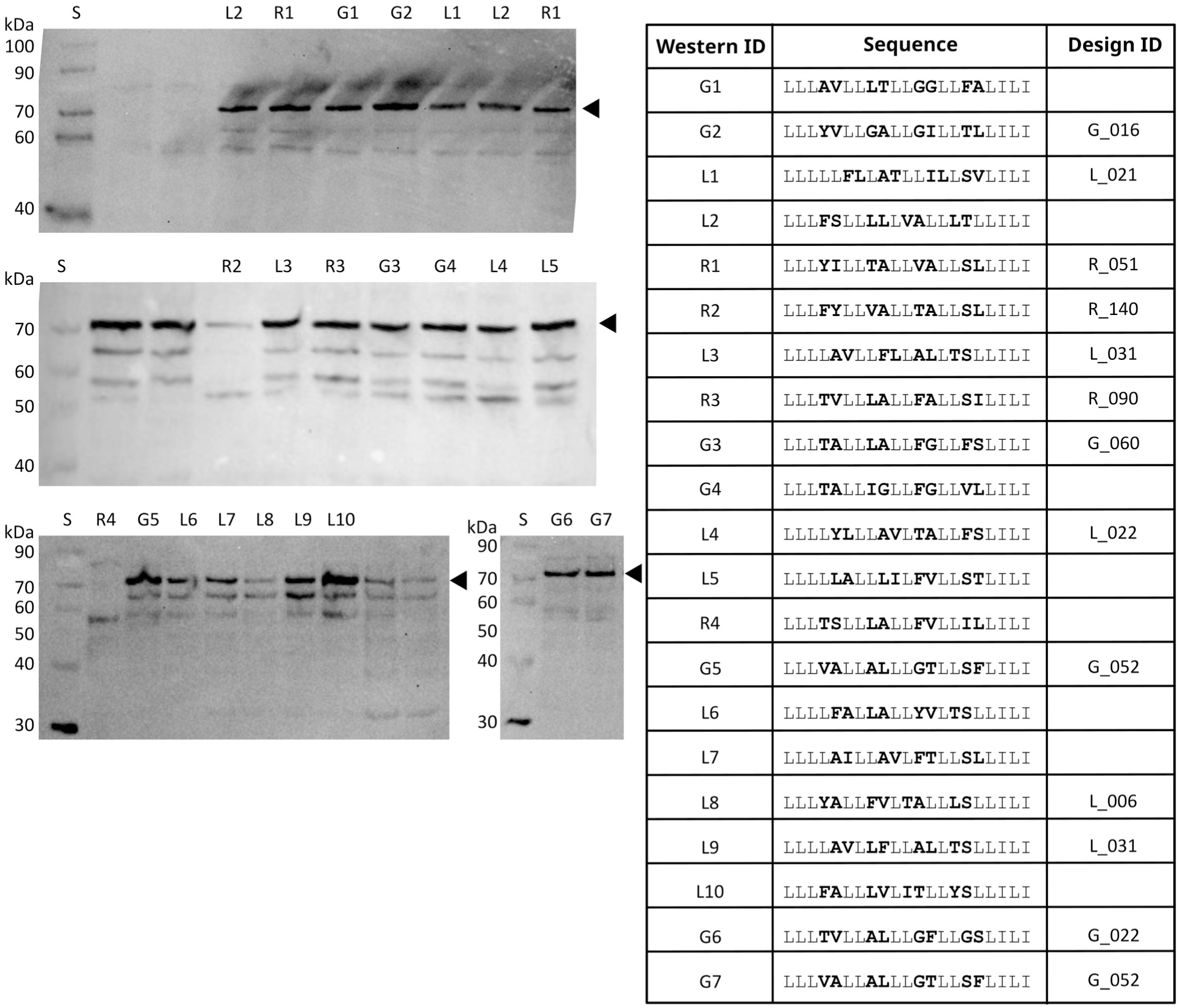
Protein expression of designs. Protein expression was confirmed by randomly sampling a number of constructs by immunoblotting with anti-maltose binding protein antibodies. L: “Left” designs. R: “Right” designs. G: GAS_right_ designs, S: molecular standards. The triangle indicates the main band of interest. Although some variability is observed (most notably R2, R4 and L10), the expression level is overall consistent across the constructs.

**Fig. S3.**
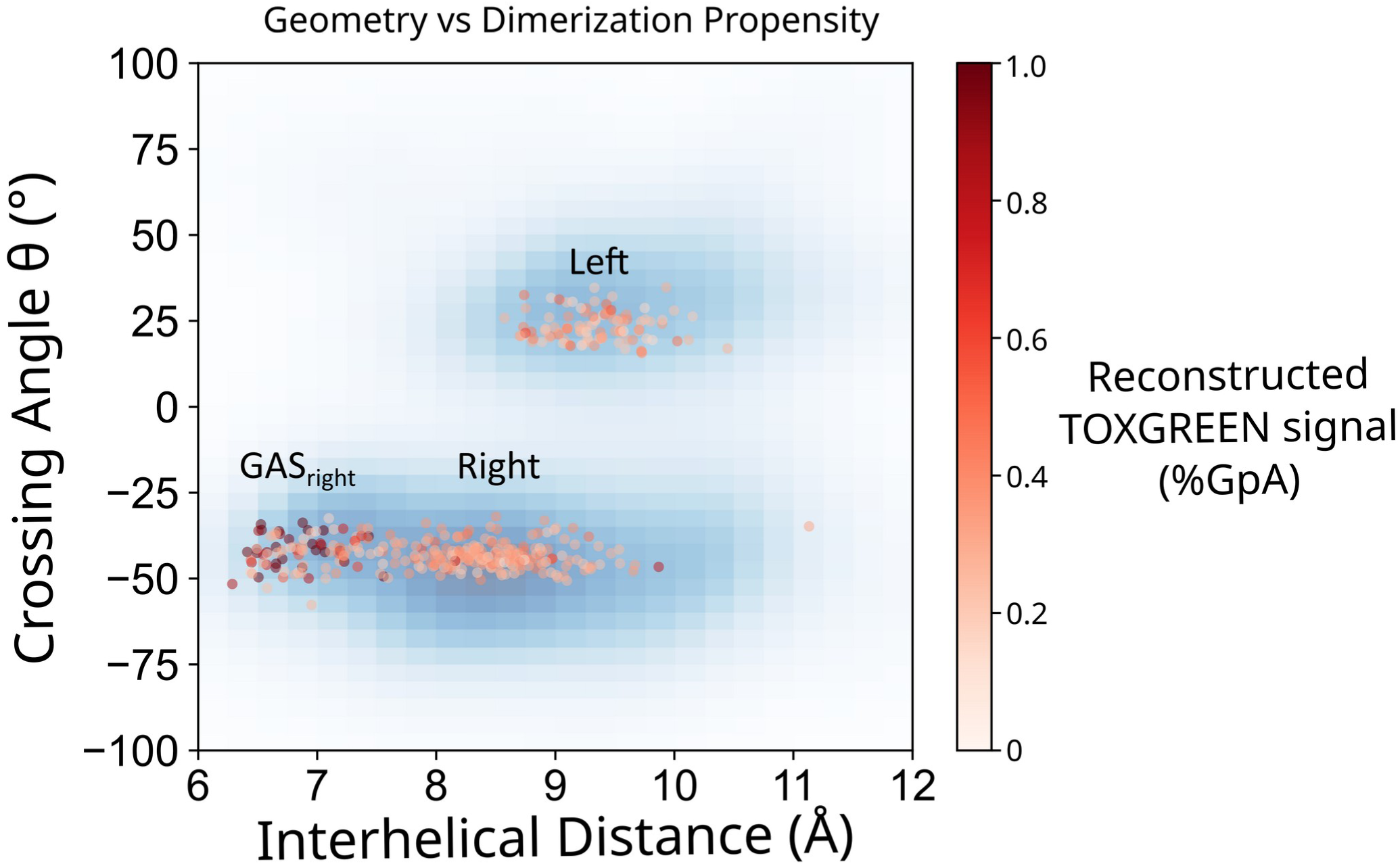
Mapping of the final set of 613 as a function of crossing angle θ and interhelical distance *d*. The color coding shows the reconstructed TOXGREEN dimerization propensity of each design.

**Fig. S4.**
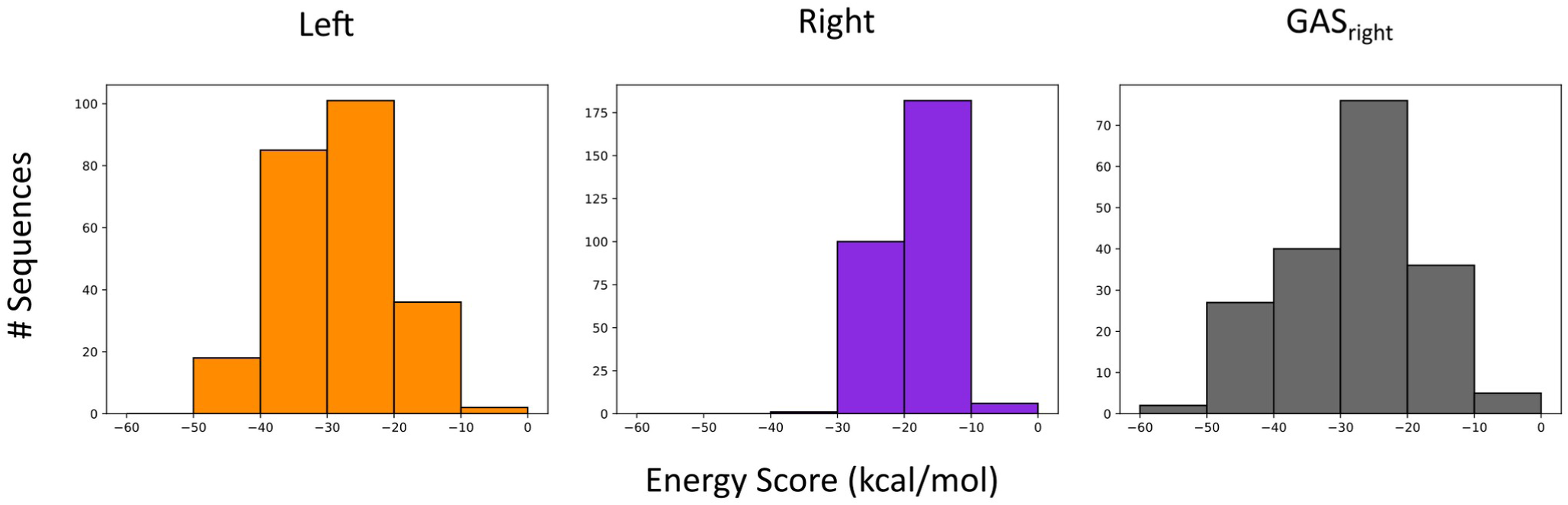
**Distribution of the computational energy score of the final set of 613 designs in the three design regions.**

**Fig. S5.**
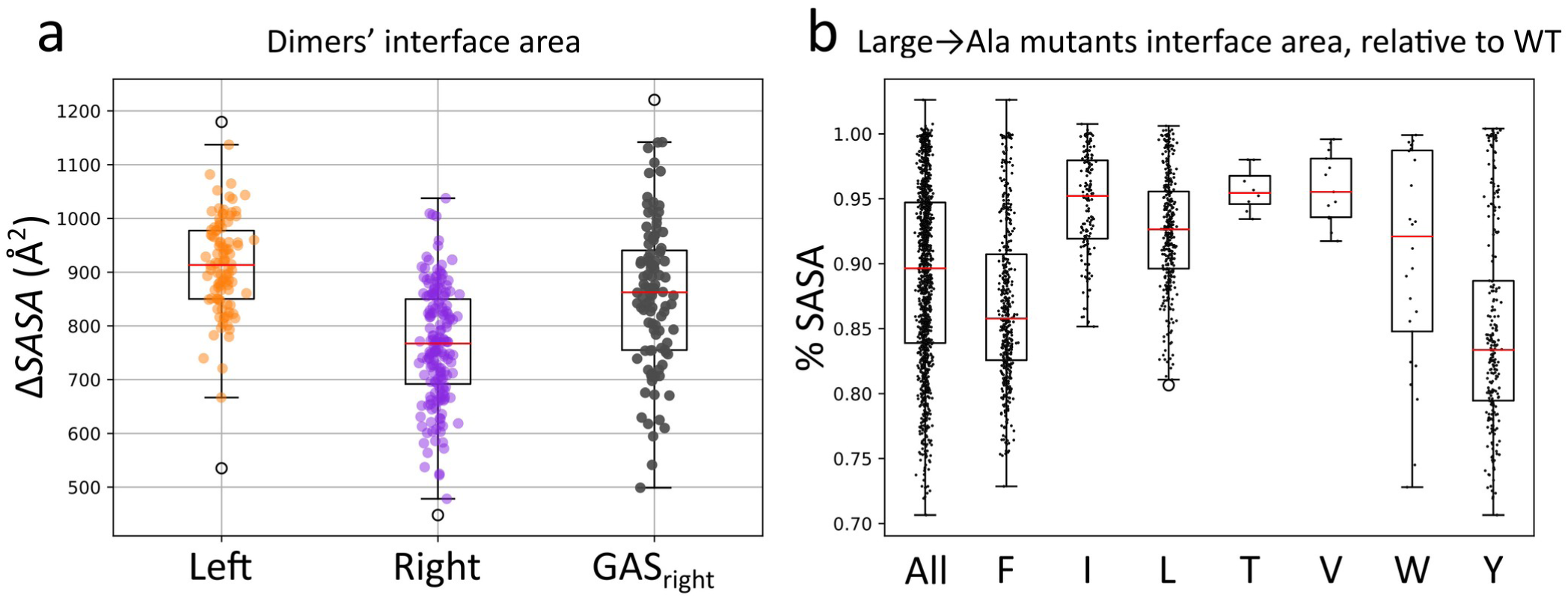
Surface area of the design’s dimer interaction interface and effect of mutation. a) Distribution of the surface area of the interaction interface in the computational models of the final set of 613 Left, Right and GAS_right_ designs. The interaction surface areas were calculated by subtracting the Solvent Accessible Surface Area (SASA) of the helix in a monomeric configuration from the SASA of the dimer. b) Change in interaction interface area of various types of mutants normalized to the area of their respective wild-type sequences.

**Fig. S6.**
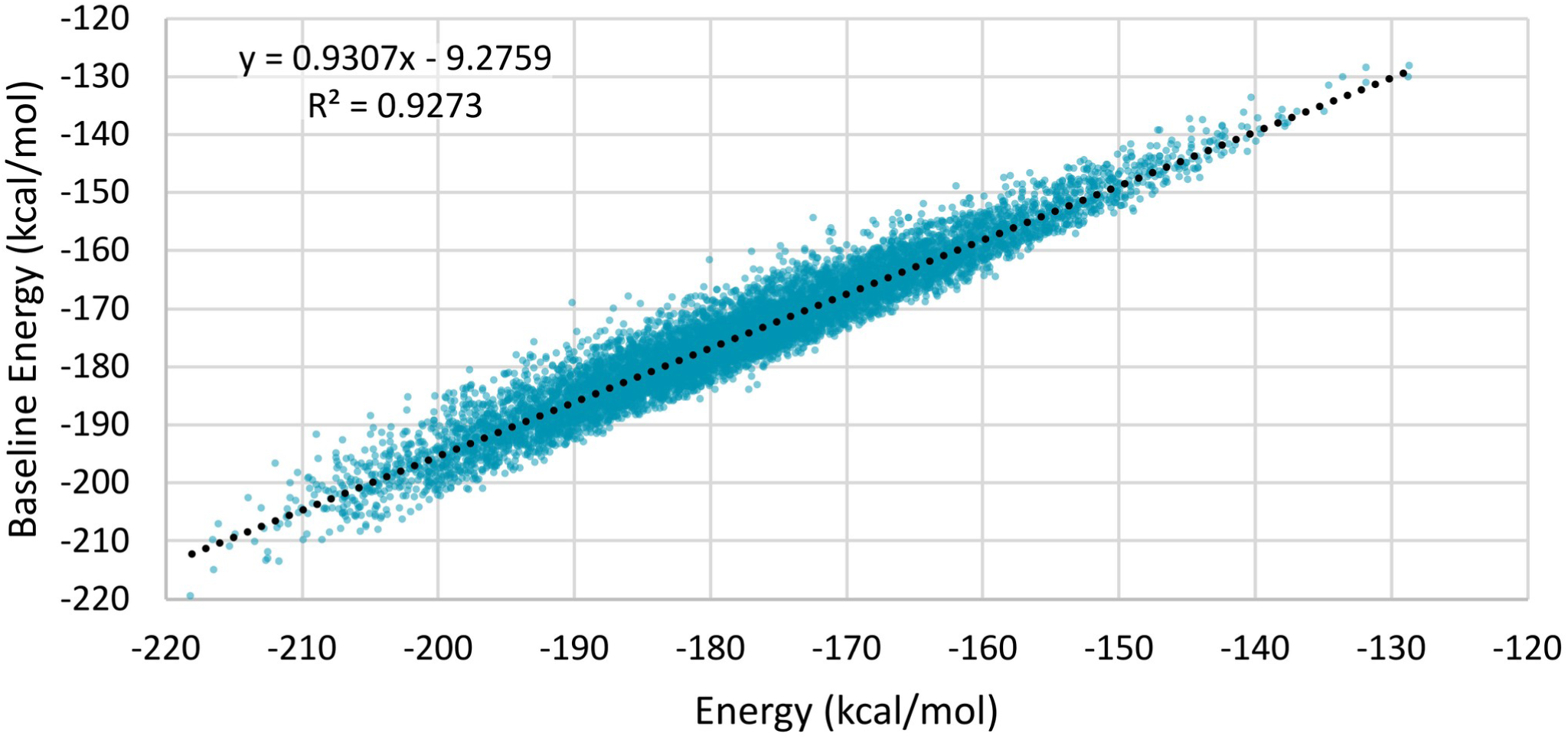
Calculation of the monomer baseline energies. A monomer energy approximation (baseline term) for approximating the energy of the monomeric helices during design. The term was developed by averaging the interactions of each individual amino acid in a large set of monomeric helices of different amino acid compositions. The baseline energies include a self term that corresponds to the interactions that occur within the single amino acid, and a pair term that corresponds to the interaction of an amino acid with other amino acids in the helix. The figure shows the comparison between the baseline energy and the actual energy computed for the corresponding monomer in 10000 random sequences in a helical configuration.

**Fig. S7.**
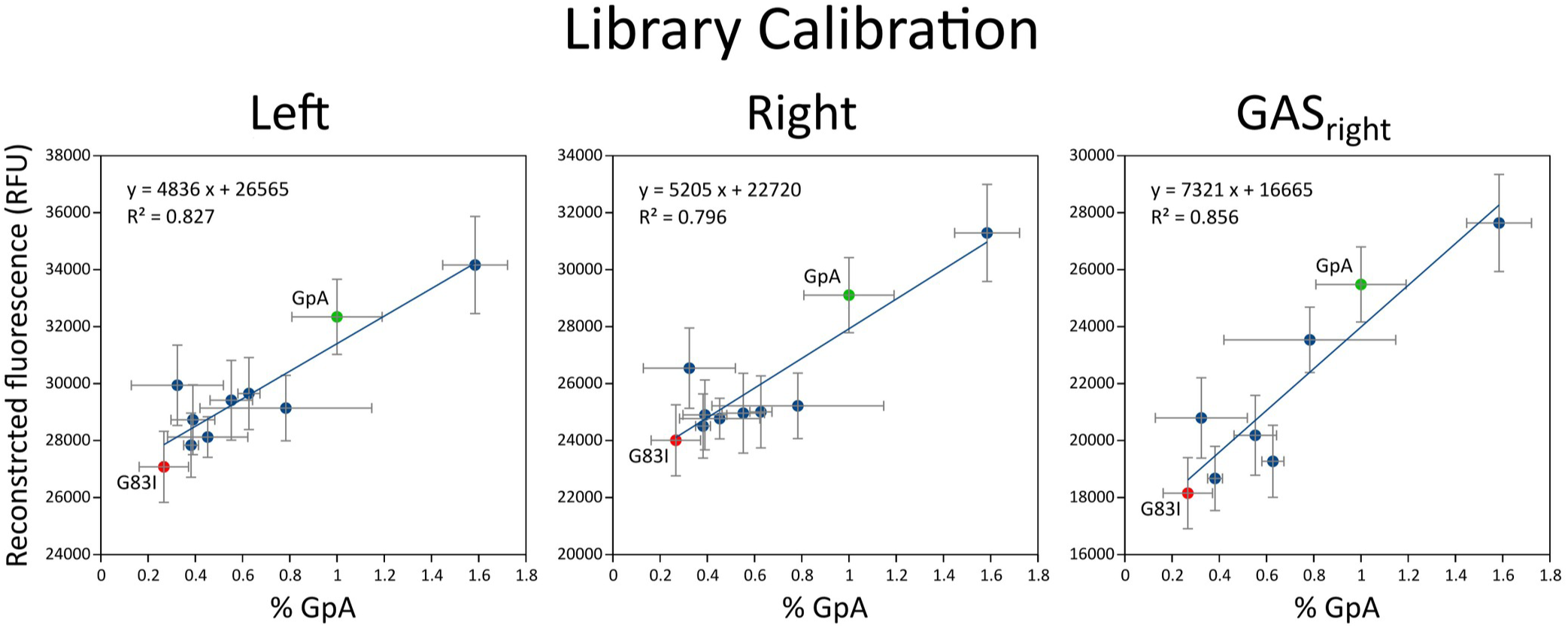
Normalization of reconstructed fluorescence to standard. The. TOXGREEN signal of a construct is typically reported normalized to the signal of the strong dimer glycophorin A (GpA, green dot). The signal is also compared to the to the monomeric variant of GpA, G83I (red dot). To avoid the dependency of the conversion on the variability of the single GpA construct, we calculated a calibration curve for each library using a set of 8-10 constructs of known TOXGREEN signal. Normalization was performed according to the resulting linear relationship. X-axis: normalized TOXGREEN signal measured individually in a fluorescence plate reader. Y-axis: reconstructed fluorescence of the same constructs in the libraries.

**Table S1.**
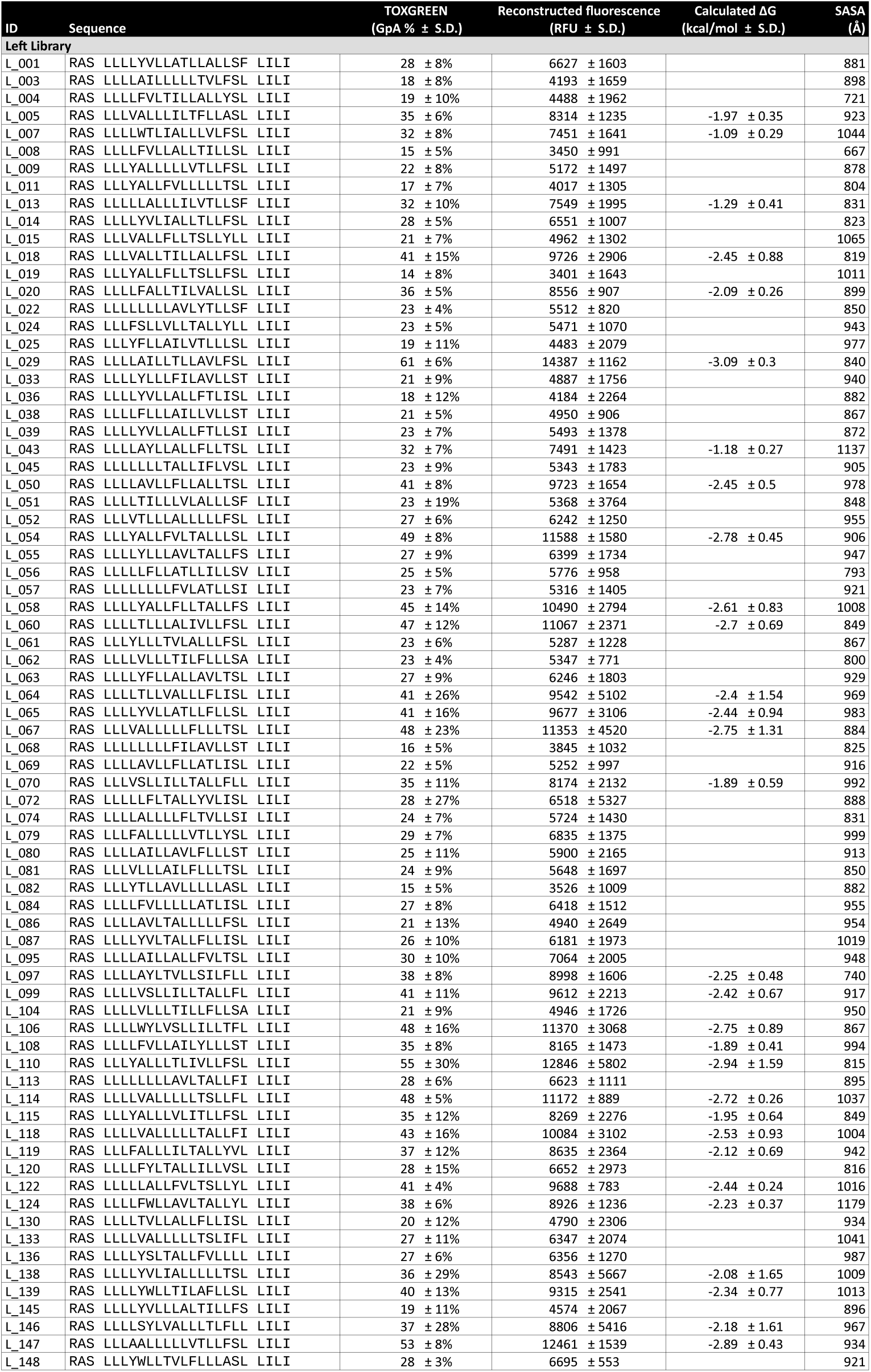
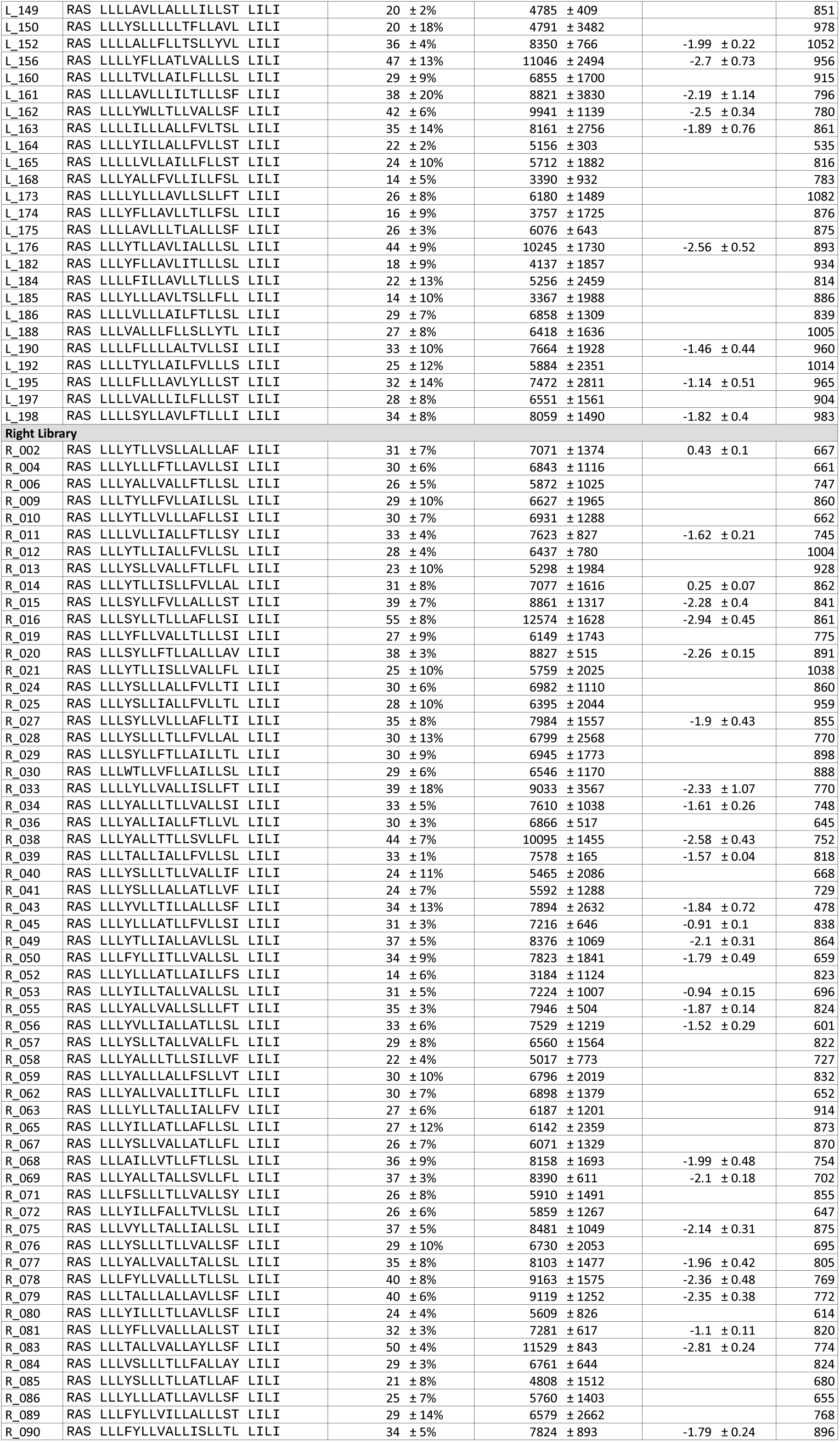
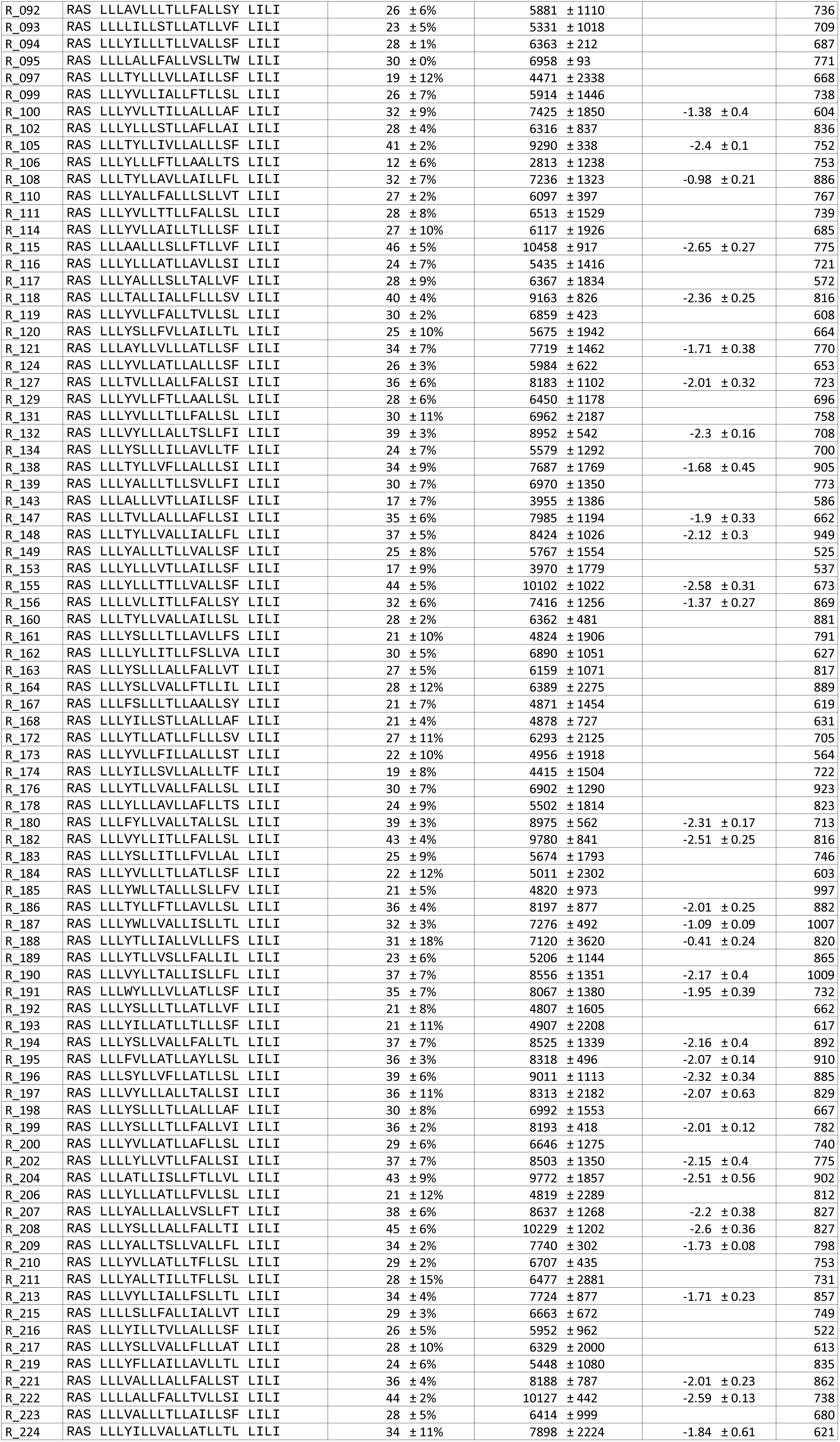
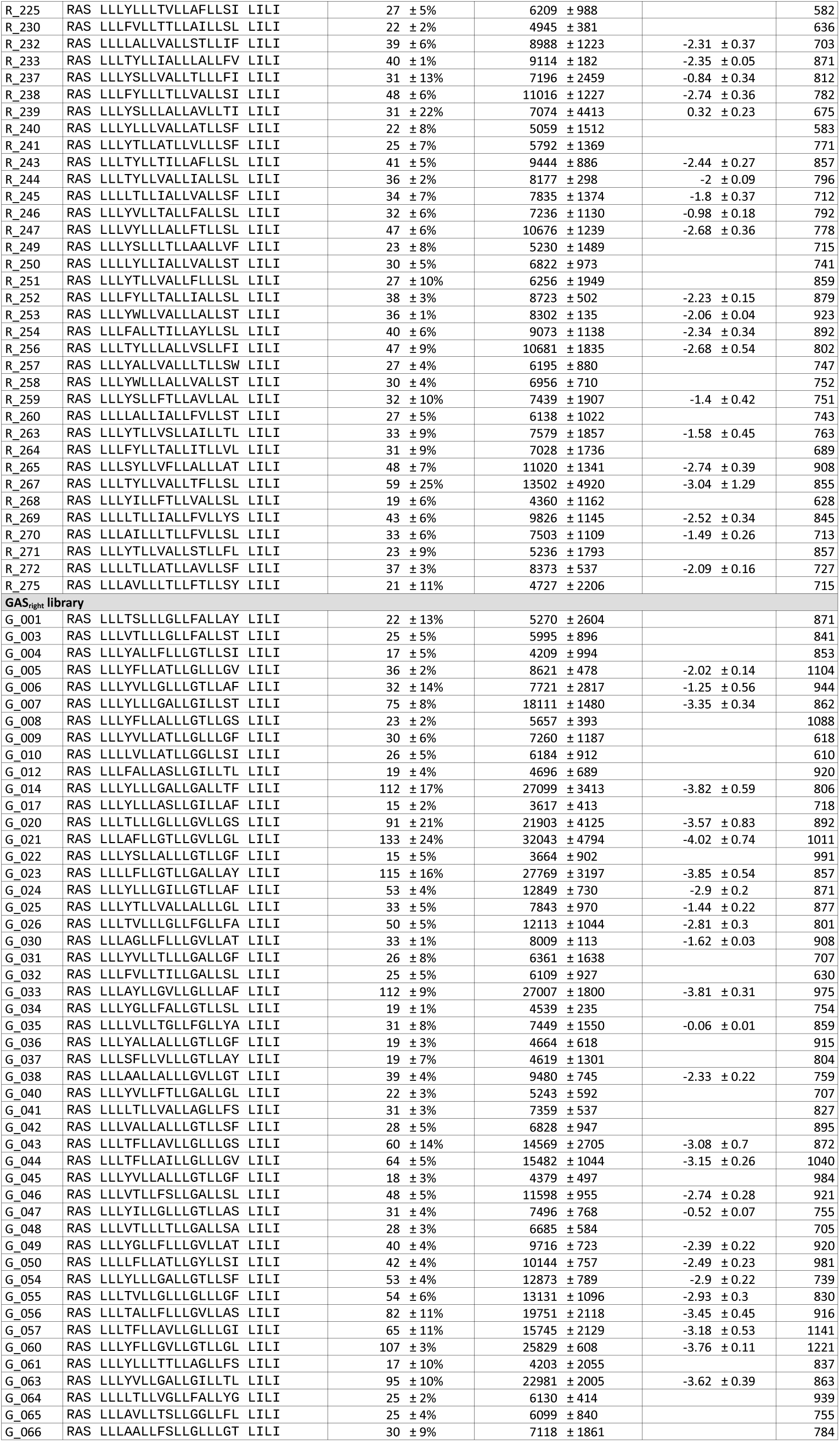
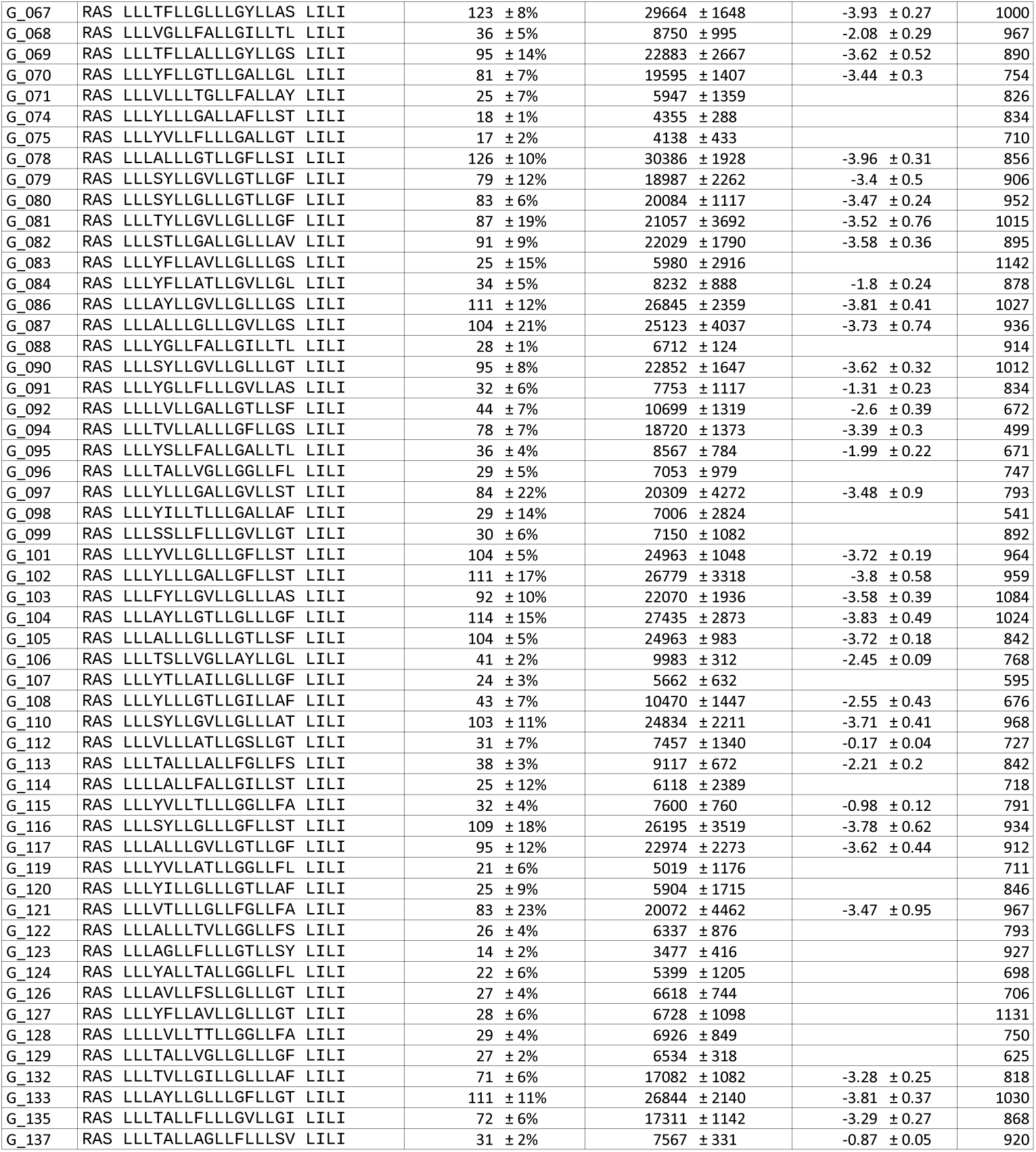
Dataset of validated designs. This dataset includes all constructs whose whose Clash mutants were in the monomeric range using >35% of the GpA TOXGREEN signal as the criterion.

**Table S2.** Frequency of the hydrophobic amino acids within the transmembrane region of membrane proteins in the OPM database.

| Amino acid | Frequency |
| --- | --- |
| ALA | 0.143 |
| PHE | 0.098 |
| GLY | 0.085 |
| ILE | 0.121 |
| LEU | 0.211 |
| SER | 0.073 |
| THR | 0.066 |
| VAL | 0.126 |
| TRP | 0.028 |
| TYR | 0.049 |

**Table S3.**
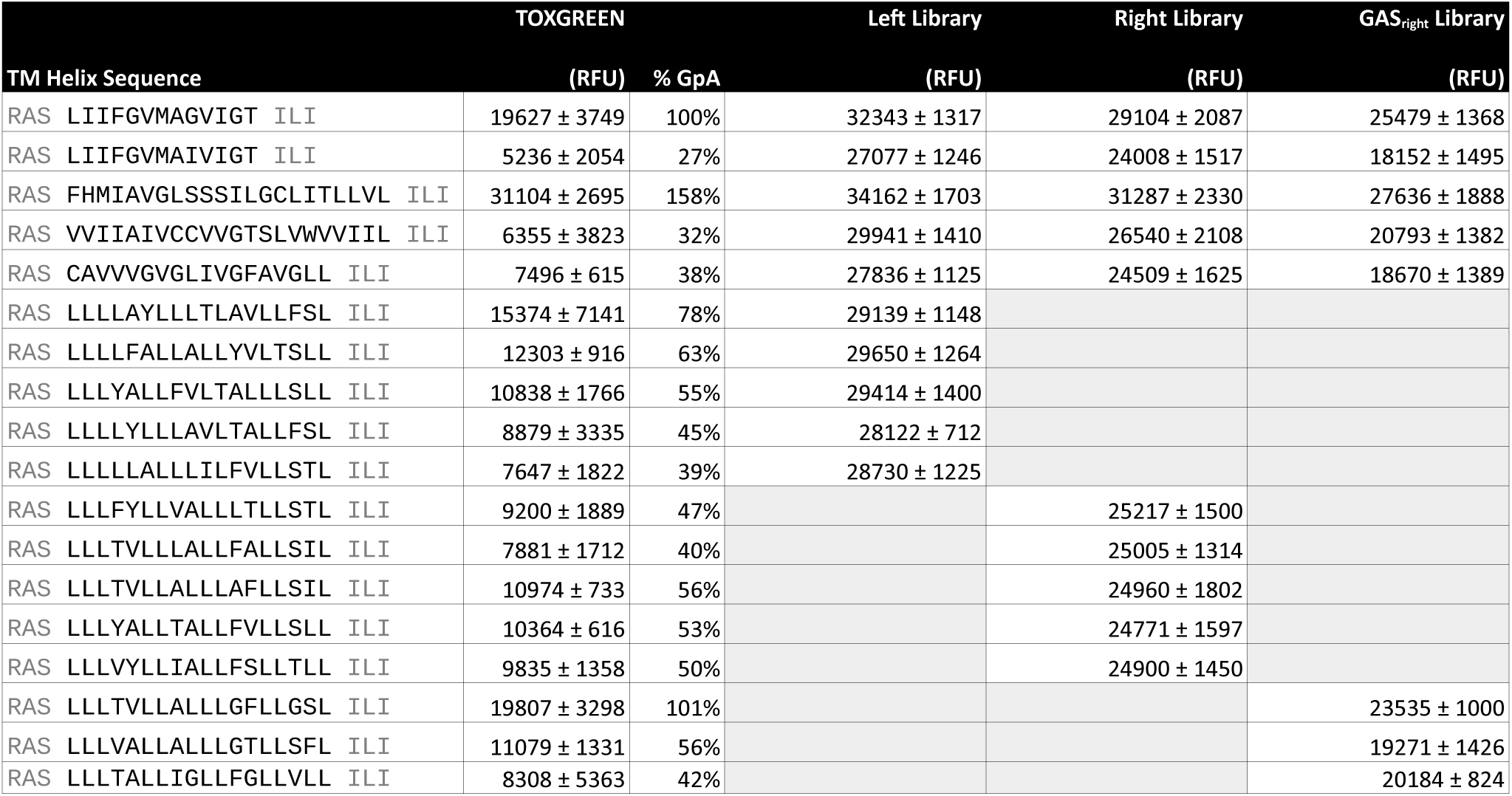
Standard constructs used for fluorescence reconstruction normalization. The table reports the individually measured GFP fluorescence (TOXGREEN) and their normalized value to the customary GpA standard (% GpA). The table also reports the reconstructed fluorescence values for the same constructs as obtained for the Left, Right and GAS_right_ libraries. Linear fitting was performed for these individual measurement and their library reconstruction vaues, as illustrated in supplementary Fig. S6. The resulting linear relationships were used used to calibrate the conversion of the reconstructed fluorescence values for three libraries to normalized % GpA values.

